# Using a micro-device with a deformable ceiling to probe stiffness heterogeneities within 3D cell aggregates

**DOI:** 10.1101/2023.10.03.559477

**Authors:** Shreyansh Jain, Hiba Belkadi, Arthur Michaut, Sébastien Sart, Jérôme Gros, Martin Genet, Charles N. Baroud

## Abstract

Recent advances in the field of mechanobiology have led to the development of methods to characterize single-cell or monolayer mechanical properties and link them to their functional behaviour. However, there remains a strong need to establish this link for three-dimensional multicellular aggregates, which better mimic tissue function. Here we present a platform to actuate and observe many such aggregates within one deformable micro-device. The platform consists of a single PDMS piece cast on a 3D-printed mold and bonded to a glass slide or coverslip. It consists of a chamber containing cell spheroids, which is adjacent to air cavities that are fluidically independent. Controlling the air pressure in these air cavities leads to a vertical displacement of the chamber’s ceiling. The device can be used in static or dynamic modes over time-scales of seconds to hours, with displacement amplitudes from a few μm to several tens of microns. Further, we show how the compression protocols can be used to obtain measurements of stiffness heterogeneities within individual co-culture spheroids, by comparing image correlations of spheroids at different levels of compression with finite element simulations. The labeling of the cells and their cytoskeleton is combined with image correlation methods to relate the structure of the co-culture spheroid with its mechanical properties at different locations. The device is compatible with various microscopy techniques, including confocal microscopy, which can be used to observe the displacements and rearrangements of single cells and neighborhoods within the aggregate. The complete experimental and imaging platform can now be used to provide multi-scale measurements that link single-cell behavior with the global mechanical response of the aggregates.

## 1. Introduction

The field of mechano-biology has experienced rapid growth in recent years, both from a fundamental science point of view [1] and for tissue engineering applications [2]. This growing interest in the link between mechanics and biology has led to the development of a wide range of techniques to measure the elastic and viscoelastic properties of individual-cells, including atomic force microscopy, optical stretchers, microfluidics, or deformable pillars [3]. As a result, the mechanical properties of cells can now be related to their state of disease [4, 5] or their differentiation potential [6], while mechanical cues are used to direct stem cell differentiation [7] or to induce an epithelial to mesenchymal transition (EMT) of cancer cells [8]. Beyond work on individual cells, collective phenomena have also been studied on 2D monolayers of cells, e.g. using traction force microscopy or image analysis [9, 10, 11].

In contrast with this highly developed field, very few methods have been able to measure the elastic or viscoelastic (the rheological) properties of 3D tissues, even though these 3D situations are more relevant to *in vivo* conditions, both biologically and mechanically. Some work has probed the forces acting within growing 3D tissues [12, 13, 14, 15], while different methods have been devised to measure the rheology of individual spheroids [16, 17, 18]. However, there is a need for a simple method that can probe the spheroid mechanobiology dynamically and generate data on multiple spheroids in parallel.

The mechanical manipulation of multiple cellular monolayers (2.5D) or 3D cellular aggregates has been performed using several microfluidic platforms, either through the stretching of a membrane [19] or the 3D compression of multicellular spheroids in a hydrogel [20]. These methods leverage the flexibility of the most common microfluidic material, polydimethylsiloxane (PDMS), to apply deformations or stresses on the biological structure. In both of these approaches, the pressure is controlled inside air-filled chambers that are positioned on the sides of the channel containing the biological material. The deformation of these chambers applies a lateral motion in the plane of the device, which is then transferred to the biological tissue through a flexible membrane [19] or a hydrogel [20]. A consequence of this choice is that the micro-fabrication requires either multi-layered devices or thin membranes that must be bonded to the glass substrate.

Here we introduce an alternative approach to studying the mechanics of 3D cultures, based on using out-of-plane deformations of the microfluidic device. The geometry, which is inspired by recent advances in the field of soft robotics [21], is simple to fabricate in a single PDMS casting step on a 3D printed mold. We show that the devices can produce well-calibrated static or dynamic loads on cell spheroids, over periods of a few seconds to several hours. Further, we also show how mechanical information about the cell state can be extracted by comparing measurements obtained from bright-field imaging with numerical simulations. Finally, we show that the device is suitable for high-resolution confocal microscopy, which opens the path to linking single-cell with tissue-level behaviors.

### 2. Micro luidic design and characterization

The microfluidic design is based on immobilizing spheroids between the roof and floor of a wide (in-plane) and flat (out-of-plane) *observation chamber* and then applying compressive forces by modulating the height of this chamber. The construction of the chamber then allows unhindered optical access to the spheroid shape and other optical measurements during the mechanical compression, as described below.

#### 2.1. Chip geometry and fabrication protocol

The micro-device is fabricated as a monolithic piece of PDMS, which is cast onto a 3D-printed mold, then bonded to a coverslip or glass slide. It is composed of two main components, as shown in Fig. 1a: two observation chambers that are surrounded by two air cavities each. The observation chambers contain the spheroids and are fluidically independent of the air cavities(see cross-section in Fig. 1b). Each of the two observation chambers is a disc of diameter 2.5 mm and height 100 μm. They are connected to each other and to the channel inlets with a 500 *μ*m high channel, as shown in the cross-section of Fig. 1c and the full scan of the device in Fig. 1e. Two air cavities surround each observation chamber. They are rectangular in the x-y plane and have a right-trapezoidal cross-section, with one rounded corner, in the x-z plane (Fig.1b). Applying a negative pressure inside the air cavities deforms the PDMS and decreases the ceiling height in the observation chamber, allowing for the compression of the spheroids.

**Figure 1.**
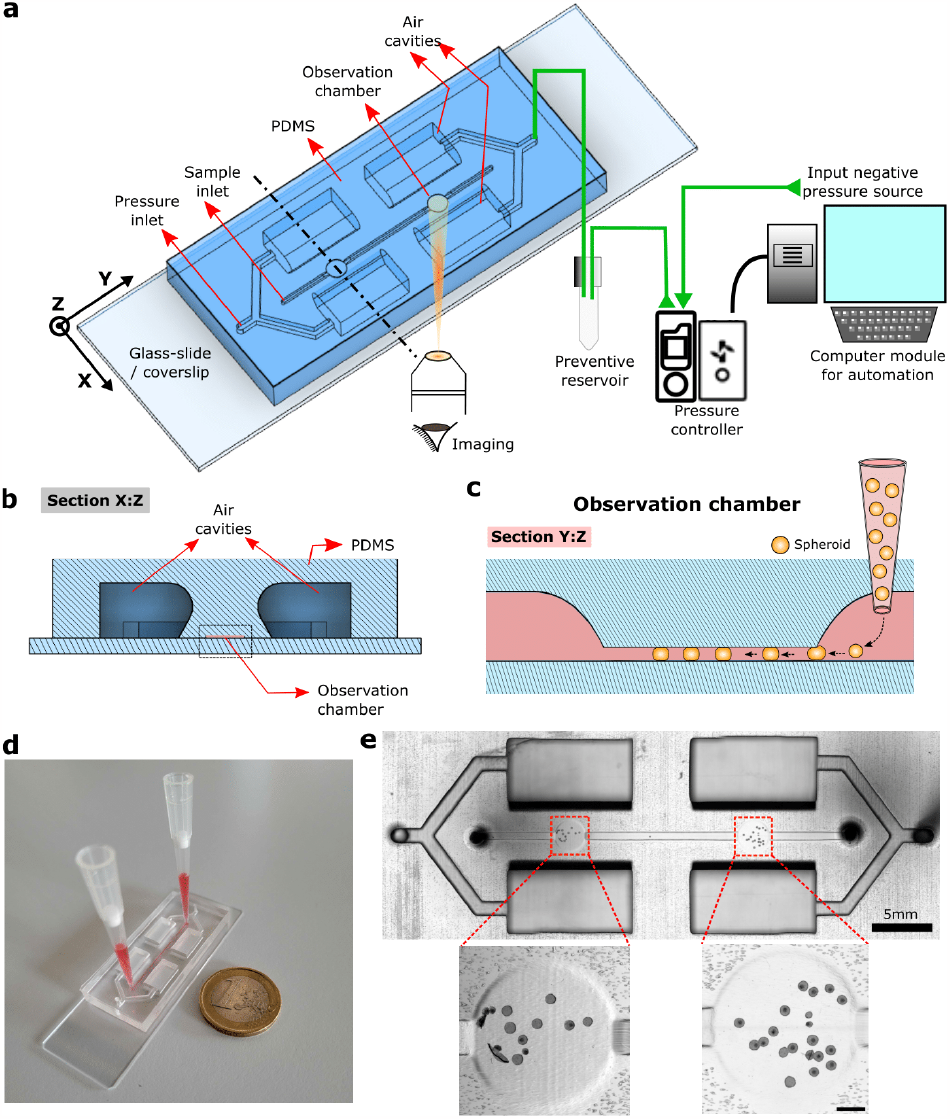
Chip design and protocol. **a**. Schematic representation of the device and the stimulation setup. The imaging is performed from the bottom of the glass slide or coverslip. **b**. X-Z Cross section of the device at the location shown by the dashed black line in panel a. The cross-sections of the air cavities and the observation chambers are the non-hatched areas. **c**. Y-Z section of the device at the location of the observation chamber disc shown in panel a. The protocol for spheroid loading is schematically represented. **d**. Picture of a chip filled with food coloring (red). **e**. Brightfield image of a chip filled with spheroids (X-Y view). Bottom image scale bar: 500*μm*.

A typical experiment begins with filling the observation chamber with the culture medium. Then preformed hepatoma and/or fibroblast spheroids (130 ±20*μ*m in diameter) are introduced at the inlet, using a hand-help pipette. The high ceiling of the channel allows them to flow freely at first. Once they reach the observation chamber they are slightly compressed, so that they become trapped by the low ceiling (Fig.1c). The spheroids can then be deformed by the moving ceiling, while their geometrical and functional responses are observed with a microscope (Fig.1e).

#### 2.2. Calibration of the ceiling deformations

Applying a negative pressure inside the air cavities deforms the device and decreases the ceiling height in the observation chamber. This deformation depends on the geometry and inner pressure of the air cavities. To control the strain applied to the spheroids, the device deformation is calibrated for different inlet pressures, as shown in Fig. 2.

**Figure 2.**
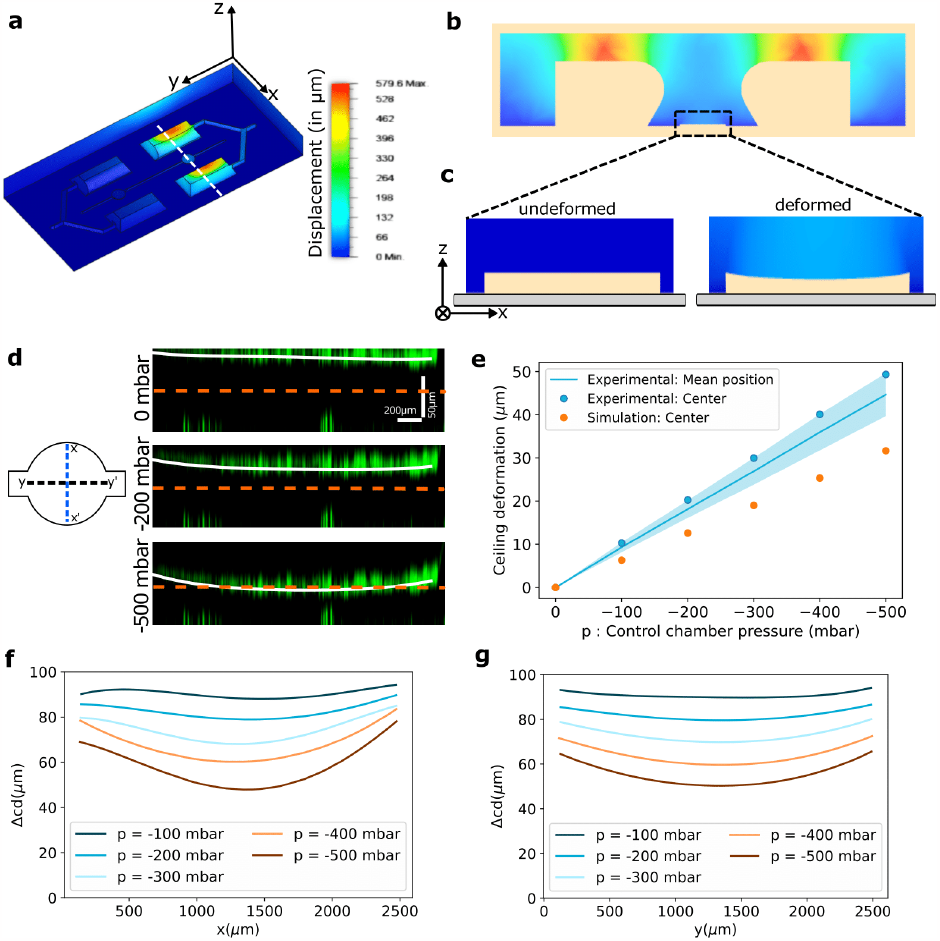
Calibrating ceiling deformation as a function of control pressure. **a** Simulation of the deformations (in *μ*m) in the PDMS device (assuming *E*_*P DMS*_ = 2 MPa, *ν*_*P DMS*_ = 0.49, *p* = −500 mbar). **b** A section in the X-Z plane of the device marked by white dashed line in panel a. **c** Magnified view from panel b for the simulation of the ceiling deformation at the observation chamber. **d** Deformation of the ceiling at the disc region of the observation chamber (along the Y direction marked by Y-Y’ in schematic) labeled with fluorescent particles present on the top and bottom surfaces of the chamber. The solid white line represents a quartic fit of the fluorescent particle positions as the ceiling goes down. **e** Ceiling deformation in the observation chamber (displacement in *μm*) as a function of the control chamber pressure. The line is the average displacement over all the positions of the ceiling. The points represent the displacement of the center (intersection of X-X’ and Y-Y’ in panel d) of the ceiling, and enable a comparison of the experiment and the simulation. **f**,**g** Ceiling deformation profile along the X-X’ and Y-Y’ represented in panel d at different pressures of the air cavities. The quartic fit of the fluorescent particles for pressure 0 mbar was subtracted from the quartic fits of the other pressure levels.

First, the device deformation was simulated using Autodesk Fusion 360 for different geometries of the air cavities and observation chambers (Fig.2a-c and SI Movie 1 for an animation). A rounded trapezoidal cross-section of the air cavities was found to induce an efficient vertical displacement of the ceiling of the observation channel, for the range of negative pressures available. In particular, the protrusion of the overhanging part above the channel could be modulated to enhance the deformation of the observation chamber. The amount of protrusion was then chosen through a compromise between increasing the deformation while maintaining the ease of microfabrication, particularly when peeling the PDMS off the 3D-printed mold.

The simulation results were then tested experimentally by injecting a suspension of fluorescent microspheres into a device and letting the solvent dry. This led the PDMS and glass surfaces inside the chip to be non-specifically coated with a layer of fluorescent particles that could be tracked by performing confocal z-stacks for different air cavity pressures, as shown in the orthogonal views along y-y’ in Fig.2d (SI Movie 2). They show that the whole ceiling indeed goes down as negative pressures are applied in the air cavities. It is important to notice that the axes in these images are scaled unequally, as is apparent from the scale bars. The ceiling is, therefore, essentially flat and curves slowly around the edges. The ceiling position is then estimated by fitting a quartic function (in white) to the positions of the fluorescent particles. Then by subtracting the fit at zero applied pressure, the vertical displacement of the ceiling could be measured at each pressure.

The experimental measurements show that the vertical displacement of the ceiling increases linearly as a function of the negative pressure, independently of the position within the chamber (Fig.2e). To compare simulation and experiments, the displacement of the chamber center is shown for each case. The simulation also predicts a linear displacement of the chamber center but underestimates its value by ≈ 1*/*3.

The profiles of the ceiling along x-x’ and y-y’ are shown in Fig.2f,g, for a moderate range of pressures. Over this range, the variability in the ceiling displacement over its surface is ≈ ±10% around the mean value. Note that it is possible to entirely collapse the ceiling by extending the pressure range.

#### 2.3. Applying static and cyclic compression on spheroids

By following the protocols above, multiple spheroids can be injected into the observation chamber with a pipette. They can then be deformed in parallel while being observed with brightfield microscopy, as shown in Fig.3a,d and supplementary movie 3 for spheroids formed from a hepatoma cell line (H4-II-EC3). Since the pressure in the air cavities is computer-controlled, it is possible to program the spheroids to be compressed either statically (Fig.3a,b) or dynamically (Fig.3c-e). As the spheroids are compressed vertically and observed from the bottom, their area increases in the brightfield images (Fig.3a). The spheroids are segmented automatically (Fig.3d), and the change in their equatorial area is monitored (Fig.3b, c, f).

In the static compression mode, the ceiling is lowered stepwise, and we wait 5-10 s for the spheroids to reach a steady state at each step before taking an image (see movie 4). The observed area of the spheroids increases as a convex function of the applied pressure, independently of the initial undeformed area within the range tested here (See Fig. 3b,c). Moreover, by normalizing the deformed area by its value at zero deformation, the curves are found to nearly collapse on a single master curve, where the ≈ 10% variability between spheroids can be attributed to differences in position within the chamber (Fig. 3c).

**Figure 3.**
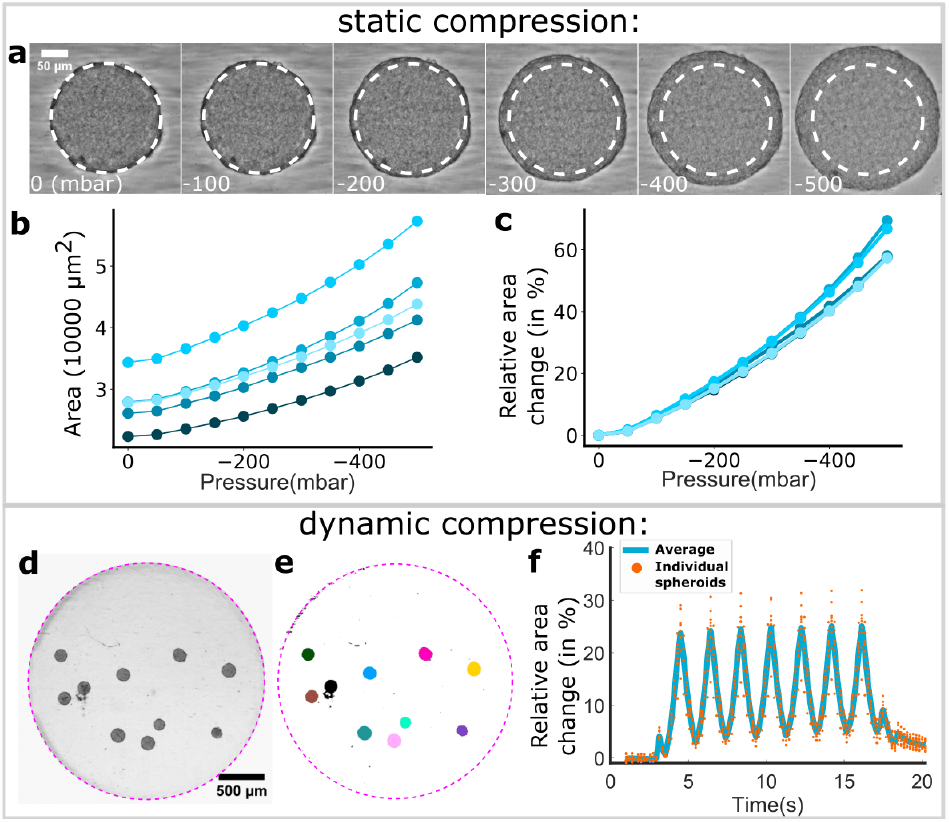
Measurements of static and dynamic spheroid compression. **a** Snapshots of a spheroid’s compression as the air cavity pressure decreases. The pressure values are on the bottom left of each picture (in mbar). **b** For 5 spheroids in the same chip (each curve is one spheroid), values of the measured equatorial area as a function of the pressure inside the control chamber. **c** For the same spheroids as b, change in the equatorial area as a function of the pressure inside the control chamber. **d** Spheroids are trapped inside the disc region of the observation chamber. **e** Automatic segmentation of the individual spheroids shown in panels c. **f** Dynamic change of spheroids’ areas (in panels c and d) during a sinusoidal pressure cycle of maximum pressure 0 mbar, minimum pressure −300 mbar, and frequency 0.5 Hz.

In addition to the static compression, the pressure air cavities can be programmed to follow a sinusoidal pattern. This in turn leads to oscillations of the chamber ceiling and of the observed area of the spheroids, as shown in Fig. 3d-f (SI Movie 3). Here oscillations in the range 0,-300 mbar, corresponding to a 25% deformation of the chamber, are applied at a frequency of 0.5 Hz. Again, the different spheroids follow the forcing frequency in synchrony, and they can be observed individually or together.

### Measuring the stiffness heterogeneity within co-culture spheroids

The ability to apply a fixed deformation of the spheroids in this device can be used to probe the mechanics of these 3D tissues, by comparing quantitative image analysis with numerical simulations, as discussed below.

#### 3.1. Biological models and their characterization off-chip

Co-culture spheroids were made by mixing H4-II-EC3 cells with fibroblasts (NIH-3T3). The result was reproducible core-shell structures where the fibroblasts formed a compact core, and the hepatoma cells formed a shell, as shown in Fig.4a. The core size was controlled with the initial number of seeded cells: NIH-3T3 cells do not proliferate in 3D while H4-II-EC3 cells do. The number of fibroblasts was therefore chosen to reach the desired core size and the number of hepatoma cells was chosen to reach a total diameter of 130 ± 20 *μ*m.

Given the different phenotypes of these cell types, we expect their mechanical properties to be different. To investigate this, we look at the structure of the cocultures by staining their nuclei and F-actin. An early indication of the mechanical contrast between the two cell types can be found by the F-actin signal, which is much brighter in the fibroblasts than in the hepatoma cells (Fig.4b) [22]. Moreover, the nuclear staining also displays a flattening of the fibroblast nuclei at the edge of the NIH-3T3 region, also suggesting that internal stresses are compressing these cells together.

To confirm that the core and shell have different elastic moduli, we made mono-culture spheroids with the two cell types and measured their elastic response in a cantilever experiment: a micro-cantilever of known stiffness was used to press the spheroid against a 3D-printed trap while observing the spheroid in bright-field (see Fig. 4c). The difference between the imposed displacement on the root of the cantilever and the observed displacement of its tip was used to measure the deflection. This allowed us to acquire the stress-strain plots of Fig. 4d, which yield average apparent Young’s moduli of *E*_*H*4_ = 2.14 ± 0.69*kPa* and *E*_3*T* 3_ = 31.8 ± 9.08*kPa*, for the hepatomas and fibroblasts respectively.

**Figure 4.**
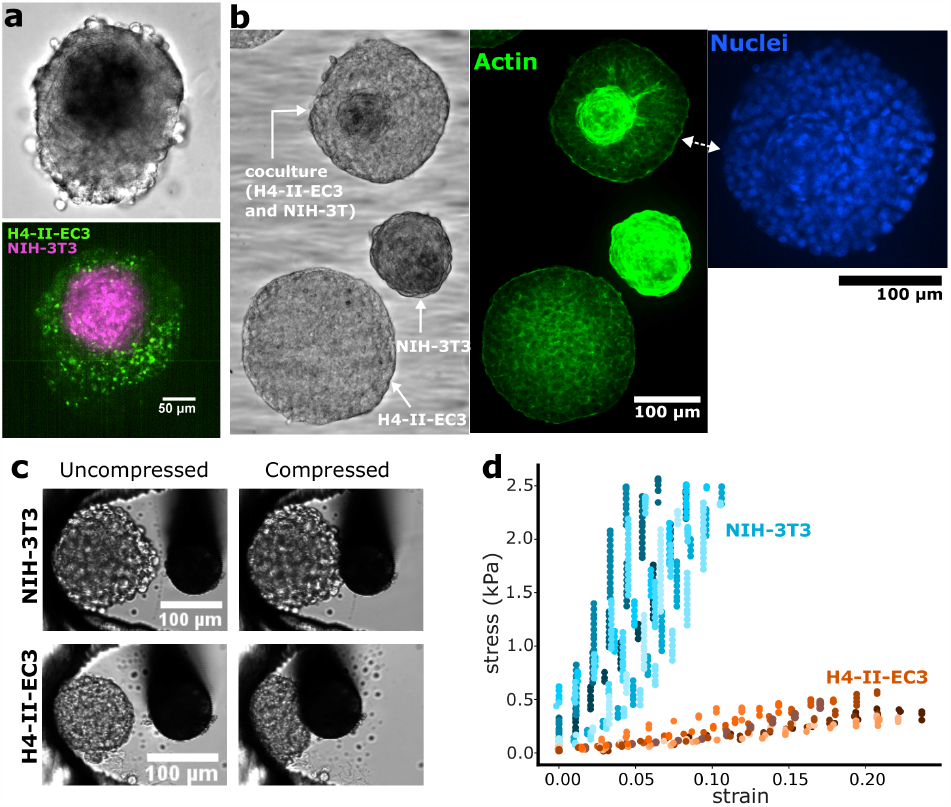
Structurally heterogeneous spheroids. **a** Vybrant dye stained coculture of NIH-3T3 (magenta) and H4-II-EC3 (green) cells. The NIH-3T3 cells form a core, and the H4-II-EC3 cells a surrounding shell. **b** Composite picture taken of three types of fixed spheroids in the chip: homogeneous H4-II-EC3 and NIH-3T3 spheroids, and a coculture of these two cell types. The images are, from left to right, a brightfield image, a fluorescence image of phalloidin-stained F-actin, and a fluorescence image of Hoechst-stained nuclei. **c** Brightfield images of homogeneous NIH-3T3 and H4-II-EC3 spheroids before (left), and after (right) compression with a cantilever. The cantilever is displaced from right to left, and the spheroids are pressed against 3D-printed traps. Starting from the moment of contact between the cantilever and the spheroid, the same displacement was imposed on the cantilever for the two spheroids in these images. **d** Stress-strain plots for multiple experiments as shown in panel c.

#### 3.2. Signature of the stiffness heterogeneity in the microfluidic device

The deformation of the monoculture and co-culture spheroids were imaged using the micro-device while applying air cavity pressures from 0 to −300 mbar, with steps of −50 mbar. Brightfield images were taken while statically compressing mono-culture hepatoma spheroids and co-cultures with different fibroblast core sizes (Fig.5 and SI movie 4). Consecutive images were compared together by applying a local image correlation algorithm, using a library for particle image velocimetry (PIV, see Methods section). This analysis provided a local displacement field on the spheroids, as shown in Fig. 5i for the transition from −100 to −150 mbar for three co-culture configurations. The norm of the velocity vectors from this vector field can then be averaged azimuthally to obtain a one-dimensional representation of the displacement as a function of the distance from the center of each spheroid. This is shown in Fig. 5iii for the three spheroids in part ii. A strong contrast is observed between the displacement field of the monoculture spheroid and the co-culture spheroids: In the first case, the displacement is continuous from the center to the edge of the spheroid, while the co-culture cases display a flat region, with very little displacement. The undeformed region matches the

**Figure 5.**
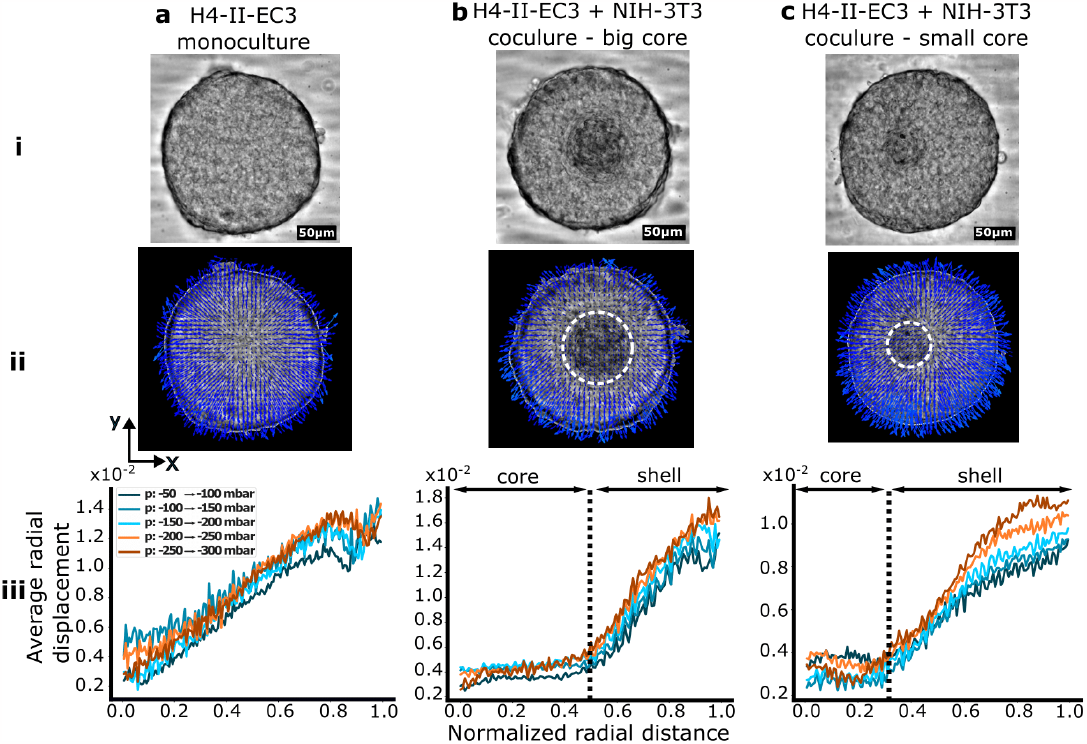
Mean-field deformations for mono-culture and co-culture spheroids. **a** Mono-culture H4-II-EC3 spheroid, **b** H4-II-EC3 and NIH-3T3 co-culture with a large core, **c** H4-II-EC3 and NIH-3T3 co-culture with a small core. **i** Bright-field images of the spheroids. **ii** PIV between one compression level and the next (with steps of −50 mbar). The dashed white lines circle the cores in the cocultures. **iii** Average radial displacement of the spheroids as a function of the position along the normalized equatorial radius (average norm of the PIV vectors in thin rings of increasing size). The dashed lines represent the core’s relative radius (respectively 0.51 (b) and 0.33 (c) of the total spheroid radius)

#### 3.3. Comparison with finite element simulations of a linear elastic model

Finite element simulations were used to better understand how differences between the mechanical responses of the core and the shell would influence the deformations of the whole co-cultures. Here the spheroids were modeled as nearly incompressible (Poisson ratio 0.49) elastic balls, deformed between two rigid surfaces. Because of invariance properties, the problem depends only on one spatial variable, here we choose the spheroid initial radius R, which we set to 1, and normalize all other quantities with respect to it. By using the problem symmetries, this could be reduced to simulating an axisymmetric two-dimensional section of a half hemisphere (Fig. 6a). In the simulated spheroid, the core and the shell were allowed to have different Young moduli *E*_*core*_ and *E*_*shell*_. Simulations were performed while varying *E*_*r*_ = *E*_*core*_*/E*_*shell*_ and simulating the mesh’s compression along *z*_*s*_ (Fig 6b). Again, because of invariance properties, the problem only depends on this ratio, so we fix *E*_*core*_ to an arbitrary value and vary *E*_*shell*_.

**Figure 6.**
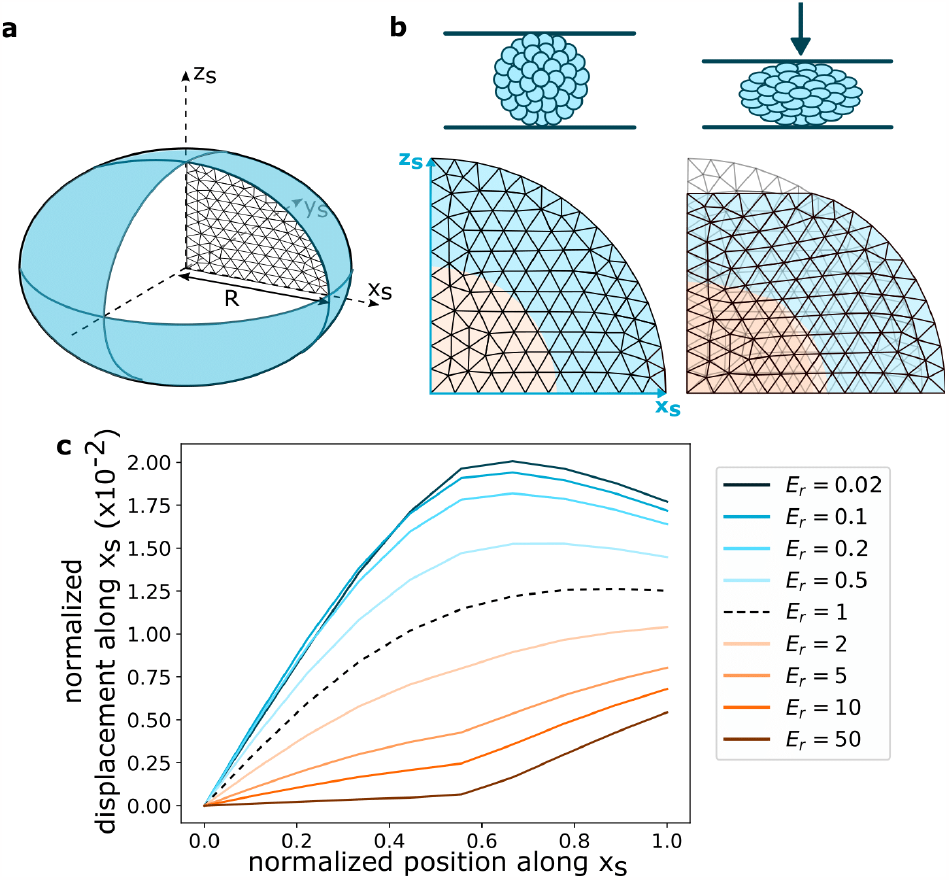
Finite-element simulations of the deformation of a spheroid. **a** Cross-section scheme of a spheroid. The simulations are performed on a half hemisphere 2D mesh of unit radius R. **b** A 2D mesh before (left) and after (right) a simulated compression. A mesh has a core, of half-unit radius (in orange), and a shell (in blue) with different Young moduli. **c** The normalized displacement along the normalized equatorial radius *x*_*s*_ for different values of *E*_*r*_ = *E*_*core*_*/E*_*shell*_ and a compression of 10% R

The displacement of the nodes along the *x*_*s*_ axis is plotted in Fig. 6c for a compressive strain of 0.1*R*. This strain is comparable with the data of Fig. 5, where the deformation due to a 50 mbar pressure change is equivalent to approximately 5 μm out of a channel height of 100 μm (Fig. 5iii). The simulations produced a family of curves for different ratios *E*_*r*_ of the core to shell moduli, as shown in Fig. 6c.

Comparison between the experimental and simulated displacement curves display semi-quantitative agreement for both the monoculture and co-culture cases. In the case of a homogeneous spheroid, the curve for *E*_*r*_ = 1 presents a concave shape that reaches a displacement at the edge of around 1.2% · *R*, which is in agreement with the experimental measurements. In the case of the co-cultures, the shape of the experimental displacement curve is consistent with a large heterogeneity in stiffness between the core and shell (*E*_*r*_ *>* 10), which is in agreement with the independent measurements using the cantilever that gives *E*_*r,exp*_ ≈ 15. Note, however, that the PIV measurements are sensitive to image elements out of the mid-plane of the spheroids. Consequently, the comparison with the simulated curves is not expected to yield a fully quantitative agreement.

## Discussion and outlook

In recent years the field of soft robotics has shown how the geometry of 3D structures could be leveraged to produce large and controlled deformations [23], namely by using pressurized channels to induce large deformations in elastomers [21]. Here we show that similar approaches of using pressure to deform an elastomeric micro-device can apply fine spatial and temporal control on the scale of a cell spheroid. Indeed, the ability of 3D printers to produce features with a large size contrast and with complex 3D shapes provides a unique opportunity to design micro-devices whose deformations can be tuned to a wide range of applications. Moreover, from a practical point of view, the 3D printing step allows design iterations to be made easily and enables different shapes to be tested. Finally, coupling this device with a programmable pressure source allows deformation to be controlled to the micron-scale and with the ability to reach frequencies that are much faster than biological times.

The device can be used to manipulate multicellular spheroids in several ways. The above measurements of mechanical heterogeneity within co-culture spheroids show that it is possible to obtain quantitative biophysical measurements from simple experimental protocols. Here, the combination of fluorescent staining of the cells and of the actin fibers was combined with the mechanical characterization of the spheroids to yield semi-quantitative measurements of the stiffness landscape within each spheroid. The resulting mechanical properties that are observed emerge from the organization of the cells in the 3D structures. While they depend on the phenotype of the different cell types, they are only indirectly linked to the stiffness of individual cells in isolation. As such, the current measurements highlight the importance of working with multicellular structures rather than only with individual cells.

While we studied here a simple core-shell structure, future work will address more complex tissues, including stem-cell derived organoids that contain a wide diversity of cell types and where the cell patterning reflects their biological state [24]. Similar deformation analysis on complex tissues will be able to show the presence of anisotropic elastic moduli that may emerge e.g. due to the alignment of contractile muscle cells [25]. Beyond simple measurements, the device can also be filled with hydrogels in order to guide the differentiation of cells within the spheroids, as was elegantly demonstrated recently [26].

However going further into the biological questions will require the ability to link the mechanical stresses and biological response on the scale of individual cells, as well as the complete tissue. Such a level of detail is critical, since force transmission is inhomogeneous within 3D cellular structures, where individual cells may experience a wide variety of mechanical stresses under the action of external forcing. Indeed confocal images show that some cells within the aggregates get compressed while others rotate or get sheared (see Figs. 7 and SI Movie 5). The ability to perform such highly-resolved images while applying the mechanical stimuli can then be combined with advanced image analysis and graph-based representations [27] in order to build multiscale models of mechanobiology [28].

**Figure 7.**
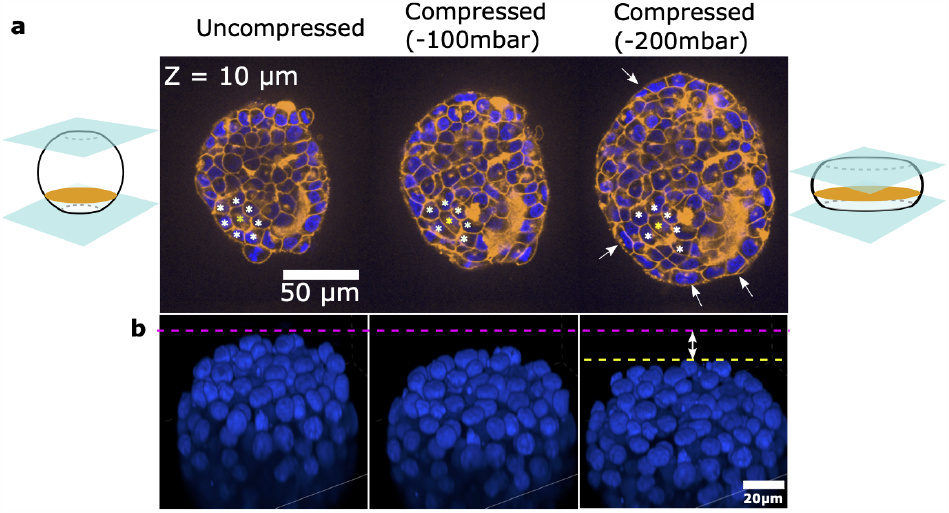
Single-cell characterization of deformations. **a** Confocal sections of a spheroid with live membrane (orange) and nuclear (blue) stainings. The panels show a section at the same z position as the spheroid is compressed (from left to right). The yellow star tracks one cell, and the white stars its neighbors, showing a change in the structure of neighboring cells. The white arrows show the re-alignment (Tangential) of the cells along the border when the spheroid is compressed, as a consequence of the mechanical stresses. **b** 3D view of the nuclei of another spheroid as it is compressed, showing the deformation or displacement of individual nuclei.

## 5. Acknowledgements

The authors acknowledge support from the Biomaterials and Microfluidics (BMcF) and the FabLab at Institut Pasteur. Shirine Merlo is acknowledged for preliminary experiments on the deformable device. HB was partially funded by the Institut Pasteur cancer initiative. JG and AM acknowledge funding by ANR-19-CE13-0024-01.

## 6. Author contributions

SJ and CNB conceptualized the project. SJ and HB conducted the device experiments and analysis. AM established the cantilever setup and AM, HB, SJ conducted cantilever experiments & analysis. HB and MG performed numerical simulations. SJ, HB, SS, AM and CNB wrote the manuscript. All authors read and discussed the manuscript.

## 8. Data Availability

The 3D print design file of the device is available on GitHub https://github.com/BaroudLab/MechanoChip. All the other raw data will be available upon request.

## 9. Materials and Methods

### 9.1. Fabrication of the compression device and setup

The molds to fabricate the chips were designed using Fusion 360 (Autodesk) and 3D printed using an SLA-FORM3 printer and ClearV4 resin (Formlabs). The molds were filled with a mixture of PDMS (SYLGARD, Dow) base and a curing agent at a ratio of 1:10. The PDMS was cured at 80°C for 2 hours. Then, it was separated from the molds and then plasma treated (Cute, Femto Science Inc.) for 40 seconds. It was then bonded to a 24 × 65 mm (#1.5) coverslip (Menzel-Gläser). The device was connected to Fluigent’s Flow EZ pressure controller (LU-FEZ-N800), fed by a supply line of −800 mbar. The pressure cycles were controlled using Fluigent’s OxyGEN software

### 9.2. Cell culture and spheroid formation

H4-II-EC3 (CRL-1600, American Type Culture Collection) and NIH-3T3 cells (CRL-1658, American Type Culture Collection) between Passage 10 and Passage 20 were maintained on T-25 cm^2^ flasks (Corning, France) in a standard CO_2_ incubator (Thermo Fisher Scientific), following the instructions provided by the manufacturer (ATCC). The culture medium was composed of Dulbecco’s Modified Eagle’s medium (DMEM) containing high glucose (Gibco, Life Technologies) supplemented with 10% (v/v) fetal bovine serum (Gibco) and 1% (v/v) penicillin-streptomycin (Gibco). The cells were seeded at 5×10^4^ cells.cm^−2^ and sub-cultivated every three days.

The spheroids were fabricated using ultra-low adhesion U-bottom 96 well plates (Corning). Monoculture of 300 H4-II-EC3 cells or 500 NIH-3T3 cells were seeded into the wells in order to form spheroids of about 130 ± 20 *μ*m in diameter after 24 hours. To fabricate co-culture spheroids, the two cell types were mixed at different ratios to generate various sizes of fibroblasts core. 100 H4-II-EC3 and 400 NIH-3T3 cells resulted in large cores, while 200 cells of each cell type gave small cores. Co-cultures were grown for 72 hours to ensure their proper core-shell arrangement by cellular self-organization.

### 9.3. Microscopy

All the images were taken using a motorized inverted microscope (Ti2, Eclipse, Nikon), equipped with either a spinning disc module (W1, Yokogawa) or an epifluorescence setup (Lumencor). Brightfield and fluorescent images were taken with a 20x objective with a 1.8-mm working distance (long working distance) and a 0.70 numerical aperture (NA) (CFI S Plan Fluor LWD, Nikon).

### 9.4. F-actin, membrane and nucleus staining

All the following reagents were introduced simply by placing a filled pipette tip at an inlet and letting the solution flow by gravity. No pressure is applied through the pipette, allowing the spheroids to stay in place. For F-actin staining, the cells were first fixed with a 4% (w/v) PFA (Alpha Aesar) for 30 min and permeabilized with 0.2 to 0.5% (v/v) Triton X-100 (Sigma-Aldrich) for 5 min. The samples were then blocked with a 5% (v/v) FBS solution and incubated for 90 min in a 1:200 phalloidin–Alexa Fluor 594 (Life Technologies) diluted in a 1% (v/v) FBS solution. The cytoplasmic membranes were stained with the CellBrite® Steady 650, following the manufacturer’s protocol. The nuclei were labeled using the NucBlue™ Live ReadyProbes™ Reagent for live cells and NucBlue™ Fixed Cell ReadyProbes™ Reagent for PFA fixed cells, following the manufacturer’s instructions.

### 9.5. Image analysis

The spheroids were segmented using a custom-made FIJI macro [29]. For each pressure value, the total area, perimeter, and major and minor axes of the spheroids are measured and recorded. To quantify radial displacement within spheroids upon compression, Particle Image Velocimetry (PIV) was performed using the open-source MatPIV toolbox in Matlab (2019b)[30], with a chosen window size 32 × 32*px* and overlap 75%. A displacement field was obtained for every deformation step applied to the spheroid by integrating the velocity data. Note that Digital Image Correlation (DIC) [31, 32] could have been used as an alternative to PIV to directly extract displacement data from the images.

### 9.6. Measurement of spheroid stiffness using cantilever

#### Cantilever fabrication and calibration

The cantilever consists of an 80*μm* diameter flexible nitinol fiber (NitiWire, Fort Waynes Metals) attached to a solid L-shaped glass pipette (Sutter instrument), held in place using a pipette holder (Narishige). We calibrated the cantilever by measuring its horizontal deflection while adding small plasticine weights. A horizontally placed stereomicroscope monitored the deflection.

The computed stiffness of a cantilever of length *l*_*ref*_ = 14.5 mm is *k*_*ref*_ = 0.2378 mN*/*mm. Then, using beam theory, we can calculate the stiffness of any cantilever of a given length. The relation between the stiffness *k* and the length *l* is 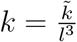, with 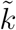 is a constant. We can then compute the constant 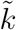 using our calibration values for *k*_*ref*_ and 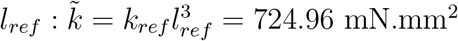.

#### Fabrication of traps for the spheroids

We fabricated traps to immobilize the spheroids during the measurement. The traps were 3D printed on coverslips using the DLP - Envisiontec Microplus HD printer. The coverslips were silanized with 3-(Trimethoxysily)propyl methacrylate to prevent the resin traps from detaching.

#### Spheroid compression

Spheroids were immobilized against 3D-printed U-shaped traps and immersed in HBSS 1X at 20°C (for less than 2 hours). They were then compressed against the traps using a vertically mounted cantilever. The cantilever was brought near the spheroid using an XYZ manual micro-manipulator (Thorlabs), coupled to a piezoelectric stage (Physik Instrumente) controlled by a computer. We imposed a linear displacement on the cantilever to compress the spheroid (a 150 μm-displacement for H4-II-EC3 spheroids and a 300 μm-displacement for NIH-3T3 spheroids). We applied a speed of 2.5 μm/sec for both cell types. We recorded the position of the cantilever and spheroid using a confocal microscope (Zeiss LSM 980) with 2-sec intervals between frames.

#### Spheroid elasticity measurement

To compute the deflection of the cantilever tip, we measured the relative motion between the piezoelectric stage and the cantilever tip. The tip position was automatically tracked using a pattern recognition algorithm (skimage.feature.match_template from the Python Scikit-Image package[**?**]). The stage position was recorded by the stage hardware. The two sets of positions were synchronized using their absolute time, and the difference between the tip and stage positions was then computed at any time by linear interpolation. The zero-deflection was defined as the offset between the tip and stage positions when the cantilever is at rest before any movement. Any deviation to this offset is then considered a deflection. To measure the strain, an ellipse was manually fit in each frame to the spheroid. The strain in each frame is defined as 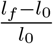, with *l* the diameter of the ellipse semi-axis along the direction of the compression, and *l*_0_ the initial value of this diameter. The strain is *f/A* with *f* the applied force and *A* the contact area of the cantilever with the spheroid. We do not know this contact area but assuming it is similar for both kinds of spheroids, we approximate *A* = 40 × 40*μm*^2^.

### 9.7. Numerical simulations

#### 9.7.1. Modeling device deformation

The ceiling’s deformation was computed by solving a standard static load application problem in Auto-desk FUSION 360. Finite element analysis of device deformation. The elastic modulus and the Poisson ratio of PDMS were input as, respectively, *E*_*PDMS*_ = 2 MPa, *ν* = 0.49. We applied negative pressure values from 0 to −500 mbar in the control chamber. The software meshes the provided geometry automatically to determine the ceiling’s deformation.

#### Modeling the mechanical response of hetero-spheroid compression

A standard linear elasticity model was solved by performing a finite element computation using the FEniCS library [33],[34] and the COMET demos [35]. Given the symmetry of spheroids and the considered compressive load, the system was reduced to an axisymmetric quarter disc section, as shown in figure 6a. Therefore, we simulated the compression along the vertical axis on a mesh of such a 2D section (Fig. 6b), and extracted the displacement of the nodes along the horizontal axis. The mesh has a core and a shell of different Young’s moduli *E*_*core*_ and *E*_*shell*_. The input parameters were each Young’s modulus, the size of the spheroid, the core radius, the Poisson ratio, the amplitude of the compression, and the node density in the mesh. Convergence tests were performed by using meshes with node distances between 0.2*R* and 0.01*R*. The node distance 0.1*R* was chosen.

## 10. Supplementary

***Movie 1*** Simulation of the platform on Autodesk Fusion 360. Negative pressure is applied in the air cavities and the colors represent the displacement of the PDMS.

***Movie 2*** Deformation of the ceiling at the disc region of the observation chamber on the y-z plane, as a function of the pressure in the air cavities. The top and bottom surfaces of the chamber are labeled with fluorescent particles.

***Movie 3*** Brightfield imaging of the cyclic compression of the spheroids in Fig. 3d,e,f.

***Movie 4*** Static compression of the three spheroids in Fig. 5.

***Movie 5*** Confocal sections of the same spheroid at three levels of compression, each frame is a section at a different z position. The spheroid’s membranes (in orange) and nuclei (in blue) are stained with live cell dyes. The different panels show different compression levels, Δ*cd* corresponds to the downward displacement of the ceiling compared to its initial position.

## References

[1] Edouard Hannezo and Carl-Philipp Heisenberg. Mechanochemical Feedback Loops in Development and Disease. Cell, 178(1):12–25, June 2019.

[2] Sudong Kim, Marina Uroz, Jennifer L. Bays, and Christopher S. Chen. Harnessing Mechanobiology for Tissue Engineering. Developmental Cell, 56(2):180–191, January 2021.

[3] Garrett F. Beeghly, Kwasi Y. Amofa, Claudia Fischbach, and Sanjay Kumar. Regulation of Tumor Invasion by the Physical Microenvironment: Lessons from Breast and Brain Cancer. Annual Review of Biomedical Engineering, 24(1):29–59, June 2022.

[4] Charlotte Alibert, Bruno Goud, and Jean-Baptiste Manneville. Are cancer cells really softer than normal cells? Biology of the Cell, 109(5):167–189, 2017. _eprint: https://onlinelibrary.wiley.com/doi/pdf/10.1111/boc.201600078.

[5] Marta Urbanska, Hector E. Muñoz, Josephine Shaw Bagnall, Oliver Otto, Scott R. Manalis, Dino Di Carlo, and Jochen Guck. A comparison of microfluidic methods for high-throughput cell deformability measurements. Nature Methods, 17(6):587–593, June 2020. Number: 6 Publisher: Nature Publishing Group.

[6] Rafael D. González-Cruz, Vera C. Fonseca, and Eric M. Darling. Cellular mechanical properties reflect the differentiation potential of adipose-derived mesenchymal stem cells. Proceedings of the National Academy of Sciences, 109(24):E1523–E1529, June 2012. Publisher: Proceedings of the National Academy of Sciences.

[7] Rowena McBeath, Dana M Pirone, Celeste M Nelson, Kiran Bhadriraju, and Christopher S Chen. Cell Shape, Cytoskeletal Tension, and RhoA Regulate Stem Cell Lineage Commitment. Developmental Cell, 6(4):483–495, April 2004.

[8] Spencer C. Wei, Laurent Fattet, Jeff H. Tsai, Yurong Guo, Vincent H. Pai, Hannah E. Majeski, Albert C. Chen, Robert L. Sah, Susan S. Taylor, Adam J. Engler, and Jing Yang. Matrix stiffness drives epithelial–mesenchymal transition and tumour metastasis through a TWIST1–G3BP2 mechanotransduction pathway. Nature Cell Biology, 17(5):678–688, May 2015. Number: 5 Publisher: Nature Publishing Group.

[9] Ernest Latorre, Sohan Kale, Laura Casares, Manuel Gómez-González, Marina Uroz, Léo Valon, Roshna V. Nair, Elena Garreta, Nuria Montserrat, Aránzazu del Campo, Benoit Ladoux, Marino Arroyo, and Xavier Trepat. Active superelasticity in three-dimensional epithelia of controlled shape. Nature, 563(7730):203–208, November 2018. Number: 7730 Publisher: Nature Publishing Group.

[10] Ana Lisica, Jonathan Fouchard, Manasi Kelkar, Tom P. J. Wyatt, Julia Duque, Anne-Betty Ndiaye, Alessandra Bonfanti, Buzz Baum, Alexandre J. Kabla, and Guillaume T. Charras. Tension at intercellular junctions is necessary for accurate orientation of cell division in the epithelium plane. Proceedings of the National Academy of Sciences, 119(49):e2201600119, December 2022. Publisher: Proceedings of the National Academy of Sciences.

[11] Shreyansh Jain, Victoire M. L. Cachoux, Gautham H. N. S. Narayana, Simon de Beco, Joseph D’Alessandro, Victor Cellerin, Tianchi Chen, Mélina L. Heuzé, Philippe Marcq, René-Marc Mège, Alexandre J. Kabla, Chwee Teck Lim, and Benoit Ladoux. The role of single-cell mechanical behaviour and polarity in driving collective cell migration. Nature Physics, 16(7):802–809, July 2020. Number: 7 Publisher: Nature Publishing Group.

[12] Otger Campàs, Tadanori Mammoto, Sean Hasso, Ralph a Sperling, Daniel O’Connell, Ashley G Bischof, Richard Maas, David a Weitz, L Mahadevan, and Donald E Ingber. Quantifying cell-generated mechanical forces within living embryonic tissues. Nature Methods, 11(2):183–189, 2013. arXiv: 1011.1669v3 ISBN: 1548-7105 (Electronic)\r1548-7091 (Linking).

[13] M. E. Dolega, M. Delarue, F. Ingremeau, J. Prost, A. Delon, and G. Cappello. Cell-like pressure sensors reveal increase of mechanical stress towards the core of multicellular spheroids under compression. Nature Communications, 8(May 2016):1–9, 2017. Publisher: Nature Publishing Group.

[14] Erfan Mohagheghian, Junyu Luo, Junjian Chen, Gaurav Chaudhary, Junwei Chen, Jian Sun, Randy H. Ewoldt, and Ning Wang. Quantifying compressive forces between living cell layers and within tissues using elastic round microgels. Nature Communications, 9(1):1878, May 2018. Number: 1 Publisher: Nature Publishing Group.

[15] Alexandre Souchaud, Arthur Boutillon, Gaëlle Charron, Atef Asnacios, Camille Noûs, Nicolas B. David, François Graner, and François Gallet. Live 3D imaging and mapping of shear stresses within tissues using incompressible elastic beads. Development, 149(4):dev199765, February 2022.

[16] Karine Guevorkian, Marie-Josée Colbert, Mélanie Durth, Sylvie Dufour, and Françoise Brochard-Wyart. Aspiration of Biological Viscoelastic Drops. Physical Review Letters, 104(21):218101, May 2010. Publisher: American Physical Society.

[17] Ruben C. Boot, Gijsje H. Koenderink, and Pouyan E. Boukany. Spheroid mechanics and implications for cell invasion. Advances in Physics: X, 6(1):1978316, January 2021. Publisher: Taylor & Francis _eprint: 10.1080/23746149.2021.1978316.

[18] Gaëtan Mary, François Mazuel, Vincent Nier, Florian Fage, Irène Nagle, Louisiane Devaud, Jean-Claude Bacri, Sophie Asnacios, Atef Asnacios, Cyprien Gay, Philippe Marcq, Claire Wilhelm, and Myriam Reffay. All-in-one rheometry and nonlinear rheology of multicellular aggregates. Physical Review E, 105(5):054407, May 2022. Publisher: American Physical Society.

[19] Dongeun Huh, Benjamin D. Matthews, Akiko Mammoto, Martín Montoya-Zavala, Hong Yuan Hsin, and Donald E. Ingber. Reconstituting Organ-Level Lung Functions on a Chip. Science, 328(5986):1662–1668, June 2010. Publisher: American Association for the Advancement of Science.

[20] Carlo Alberto Paggi, Bastien Venzac, Marcel Karperien, Jeroen C. H. Leijten, and Séverine Le Gac. Monolithic microfluidic platform for exerting gradients of compression on cell-laden hydrogels, and application to a model of the articular cartilage. Sensors and Actuators B: Chemical, 315:127917, July 2020.

[21] Emmanuel Siéfert, Etienne Reyssat, José Bico, and Benoît Roman. Bio-inspired pneumatic shape-morphing elastomers. Nature Materials, 18(1):24–28, January 2019. Number: 1 Publisher: Nature Publishing Group.

[22] Joseph H. Shawky, Uma L. Balakrishnan, Carsten Stuckenholz, and Lance A. Davidson. Multiscale analysis of architecture, cell size and the cell cortex reveals cortical F-actin density and composition are major contributors to mechanical properties during convergent extension. Development, 145(19):dev161281, 10 2018.

[23] Trevor J Jones, Etienne Jambon-Puillet, Joel Marthelot, and P-T Brun. Bubble casting soft robotics. Nature, 599(7884):229–233, 2021.

[24] Sébastien Sart, Raphaël F.-X. Tomasi, Antoine Barizien, Gabriel Amselem, Ana Cumano, and Charles N. Baroud. Mapping the structure and biological functions within mesenchymal bodies using microfluidics. Science Advances, 6(10):eaaw7853, March 2020. Publisher: American Association for the Advancement of Science Section: Research Article.

[25] Jia-Ling Ruan, Nathaniel L Tulloch, Maria V Razumova, Mark Saiget, Veronica Muskheli, Lil Pabon, Hans Reinecke, Michael Regnier, and Charles E Murry. Mechanical stress conditioning and electrical stimulation promote contractility and force maturation of induced pluripotent stem cell-derived human cardiac tissue. Circulation, 134(20):1557–1567, 2016.

[26] Carlo Alberto Paggi, Jan Hendriks, Marcel Karperien, and Séverine Le Gac. Emulating the chondrocyte microenvironment using multi-directional mechanical stimulation in a cartilage-on-chip. Lab on a Chip, 22(9):1815–1828, 2022. Publisher: Royal Society of Chemistry.

[27] Gustave Ronteix, Andrey Aristov, Valentin Bonnet, Sebastien Sart, Jeremie Sobel, Elric Esposito, and Charles N. Baroud. Griottes: a generalist tool for network generation from segmented tissue images. BMC Biology, 20(1):178, August 2022.

[28] Sébastien Sart, Raphaël F.-X. Tomasi, Gabriel Amselem, and Charles N. Baroud. Multiscale cytometry and regulation of 3D cell cultures on a chip. Nature Communications, 8(1):469, 2017. Publisher: Springer US.

[29] Johannes Schindelin, Ignacio Arganda-Carreras, Erwin Frise, Verena Kaynig, Mark Longair, Tobias Pietzsch, Stephan Preibisch, Curtis Rueden, Stephan Saalfeld, Benjamin Schmid, et al. Fiji: an open-source platform for biological-image analysis. Nature methods, 9(7):676–682, 2012.

[30] J Kristian Sveen. An introduction to MatPIV v. 1.6. page 27, 2004.

[31] François Hild and Stéphane Roux. Comparison of local and global approaches to digital image correlation. Experimental mechanics, 52(9):1503–1519, 2012.

[32] M. Genet, C. T. Stoeck, C. von Deuster, L. C. Lee, and S. Kozerke. Equilibrated warping: Finite element image registration with finite strain equilibrium gap regularization. Medical Image Analysis, 50:1–22, 2018.

[33] Anders Logg and Garth N. Wells. DOLFIN: Automated Finite Element Computing. ACM Transactions on Mathematical Software, 37(2):1–28, April 2010. arXiv:1103.6248 [cs].

[34] Anders Logg, Garth N. Wells, and Johan Hake. DOLFIN: a C++/Python finite element library. In Anders Logg, Kent-Andre Mardal, and Garth Wells, editors, Automated Solution of Differential Equations by the Finite Element Method: The FEniCS Book, Lecture Notes in Computational Science and Engineering, pages 173–225. Springer, Berlin, Heidelberg, 2012.

[35] Jeremy Bleyer. Numerical Tours of Computational Mechanics with FEniCS, June 2018.

